# The effects on neutral variability of recurrent selective sweeps and background selection

**DOI:** 10.1101/358309

**Authors:** José Luis Campos, Brian Charlesworth

## Abstract

Levels of variability and rates of adaptive evolution may be affected by hitchhiking, the effect of selection on evolution at linked sites. Hitchhiking can be caused either by selective sweeps or by background selection, involving the spread of new favorable alleles or the elimination of deleterious mutations, respectively. Recent analyses of population genomic data have fitted models where both these processes act simultaneously, in order to infer the parameters of selection. Here, we investigate the consequences of relaxing a key assumption of some of these studies – that the time occupied by a selective sweep is negligible compared with the neutral coalescent time. We derive a new expression for the expected level of neutral variability in the presence of recurrent selective sweeps and background selection. We also derive approximate integral expressions for the effects of recurrent selective sweeps. The accuracy of the theoretical predictions was tested against multilocus simulations, with selection, recombination and mutation parameters that are realistic for *Drosophila melanogaster*. In the presence of crossing over, there is approximate agreement between the theoretical and simulation results. We show that the observed relations between the rate of crossing over and the level of synonymous site diversity and rate of adaptive evolution in Drosophila are probably mainly caused by background selection, whereas selective sweeps and population size changes are needed to produce the observed distortions of the site frequency spectrum.

---

The effect of selection at a given locus on the properties of neutral variability at linked sites is a classic problem in population genetics, first studied by Sved (1968) and Ohta and Kimura (1970) in the context of associative overdominance – the apparent heterozygote advantage induced at a neutral locus by variants at linked loci that are maintained by heterozygote advantage or by mutation to partially recessive deleterious alleles. This work was followed by the paper of Maynard Smith and Haigh (1974) on the hitchhiking effect, where the spread of a favorable mutation reduces the level of neutral variability at a linked locus; this process has come to be termed a ‘selective sweep’ (Berry *et al.* 1991). It was later shown that selection against recurrent deleterious mutations also reduces neutral variability at linked sites by the hitchhiking process known as background selection (Charlesworth *et al.* 1993). A large theoretical literature on these topics has subsequently appeared, reviewed by Barton (2010), Stephan (2010), Charlesworth (2012a), Neher (2013) and Walsh and Lynch (2018, Chap. 8).

Much of the motivation for these theoretical studies came from the advent of data on genome-wide patterns of variability, which inspired attempts to infer the nature and parameters of selection from observations such as the relations between the level of synonymous sequence diversity in a gene and the local recombination rate (Begun and Aquadro 1992) and between its nonsynonymous divergence from a related species (Andolfatto 2007). Work of this type has recently been reviewed by Sella *et al.* (2009), Vitti *et al.* (2013), Booker *et al.* (2017) and Hermisson and Pennings (2017). Several recent studies have used the theory of the joint effects of recurrent selective sweeps and background selection, pioneered by Kaplan *et al.* (1989), Wiehe and Stephan (1993) and Kim and Stephan (2000), to estimate their effects on levels of neutral diversity across the genomes of multiple species (Corbett-Detig *et al.* 2015), and to infer the rates of occurrence of advantageous mutations and the strength of selection acting on them (Elyashiv *et al.* 2016; Campos *et al.* 2017). These studies all concluded that the level of variability in a species is often much smaller than would be expected in the absence of selection, even in regions with relatively high rates of genetic recombination. This reduction in variability reflects the effects of both selective sweeps (SSWs) and background selection (BGS), although the estimates of the parameters involved differ substantially among the different studies.

Several important assumptions underlie the model of recurrent sweeps used in this work. One is that the effect of BGS on the probability of fixation of a linked favorable mutation is well approximated by its effect on neutral variability at a site at the same location in the genome, which is described by a factor *B* that multiplies the value of *N_e_* for that site (Kim and Stephan 2000). An expression for *B* can be found from the standard equation for the effect of BGS in the presence of recombination (Hudson and Kaplan 1995; Nordborg *et al.* 1996), although this equation breaks down when the product of *N_e_* and the selection coefficient against deleterious mutations is of the order of 1 or less, especially when there is little or no recombination (Gordo *et al.* 2002; Kaiser and Charlesworth 2009; Good *et al.* 2014; Zhao and Charlesworth 2016). Using the formula of Kimura (1962) for an autosomal, semi-dominant mutation with selective advantage *s_A_*in homozygotes, the probability of fixation of a mutation with *N_e_s_A_*>> 1 in a randomly mating, discrete-generation population of size *N* is then *BN_e_s_A_*/*N* instead of *N_e_s_A_*/*N* (Peck 1994; Barton 1995; Stephan *et al.* 1999; Kim and Stephan 2000).

In addition, it is assumed that the time occupied by an adaptive substitution is negligible compared with the coalescent time, and that Hill-Robertson interference (Hill and Robertson 1966; Felsenstein 1974) among sweeps is absent, so that the times between successive sweeps are exponentially distributed and reductions in diversity can be predicted from the formula for a single sweep. Finally, the classic theory assumes that sweeps are ‘hard’, such that each favorable mutation originated as a single copy in the population, as opposed to ‘soft’ sweeps that arise from standing variation or from several independent mutational events in the same gene (Hermisson and Pennings 2005, 2017).

All of these assumptions can be questioned. The main purpose of this paper is to examine the accuracy of the assumptions concerning the effects of BGS, sweep duration and interference among sweeps, in the context of parameter values for BGS and SSWs that appear to be fairly realistic on the basis of inferences from a *Drosophila melanogaster* population (Campos *et al.* 2017). We chose to model a *D. melanogaster* population because this species has been the basis for much of the work on the effects of hitchhiking on natural variability. We used computer simulations of multiple loci that are subject to both BGS and SSWs, together with approximations for the effects of BGS and SSWs based on replacing summations across selected sites with integration. The results indicate that the standard coalescent approach to predicting recurrent sweep effects can underestimate their magnitude. We found only small effects of interference among sweeps, so that this discrepancy appears to be caused by the assumption that sweep duration can be neglected. To deal with this problem, we have developed a modified approach to predicting pairwise neutral nucleotide diversity under recurrent selective sweeps. We consider only hard sweeps, because these are amenable to simple analytic modeling and simulation. We hope to extend the models to soft sweeps in future work.

## Material and Methods

We used the simulation package SLiM (Messer 2013), version 1.8. The details of the simulation methods are described in the online manual (benhaller.com/slim/SLiM.18_manual.pdf). We modeled sets of *n* genes separated by 2kb of selectively neutral intergenic sequence (Figure 1), with all UTR sites and 70% of nonsynonymous (NS) sites subject to selection (the same selection parameters were applied to 5’and 3’ UTRs). The gene structure was chosen to represent a typical *D. melanogaster* gene (Campos *et al.* 2017). In order to simulate realistic parameters of selection, mutation and recombination for a model autosome, we rescaled the values applicable to a natural population of *D. melanogaster* by multiplying them by the ratio of *N_e_* for the population to the number of breeding individuals used in the simulations, *N,* which was usually set to 2500 (see Table 1). This conserves the products of *N_e_* and the basic parameters of selection, recombination and mutation, which control most aspects of evolution in finite populations if time is rescaled by a factor of *N*/*N_e_* (Ewens 2004).

**Table 1:** Parameters used in the simulations.

|  | **Natural population** |  | **Simulations** |  |
| --- | --- | --- | --- | --- |
| **Parameter** | **A** | **X** | **A** | **X** |
| Population size ($N$) | $1.33 \times 10^6$ | $0.997 \times 10^6$ | 2500 | 2500 |
| Rescaling factor | - |  | 532 | 532 |
| Standard effective crossover rate | $1 \times 10^{-8}$ | $1.33 \times 10^{-8}$ | $5.32 \times 10^{-6}$ | $5.32 \times 10^{-6}$ |
| G.c. rate of initiation | $1 \times 10^{-8}$ | $1.33 \times 10^{-8}$ | $5.32 \times 10^{-6}$ | $5.32 \times 10^{-6}$ |
| G.c. tract length | 440 bp | 440bp | 440 bp | 440 bp |
| Mutation rate per bp | $4.5 \times 10^{-9}$ | $4.5 \times 10^{-9}$ | $2.39 \times 10^{-6}$ | $1.79 \times 10^{-6}$ |
| Scaled selection coefficient for deleterious NS mutations ($\gamma_{NS}$) | 2000 | 20000 | 2000 | 2000 |
| Scaled selection coefficient for deleterious UTR mutations ($\gamma_{UTR}$) | 110 | 110 | 110 | 110 |
| Scaled selection coefficient for favorable NS mutations ($\gamma_a$) | 250 | 375 or 500 | 250 | 375 or 500 |
| Scaled selection coefficient for favorable UTR mutations ($\gamma_u$) | 213 | 319.5 or 426 | 213 | 319.5 or 426 |
| Proportion of NS mutations that are favorable ($p_a$) | $2.21 \times 10^{-4}$ | $2.21 \times 10^{-4}$ | $2.21 \times 10^{-4}$ | $2.21 \times 10^{-4}$ |
| Proportion of UTR mutations that are favorable ($p_u$) | $9.04 \times 10^{-4}$ | $9.04 \times 10^{-4}$ | $9.04 \times 10^{-4}$ | $9.04 \times 10^{-4}$ |
| Prop. of neutral exonic mutations | 0.3 | 0.3 | 0.3 | 0.3 |
| Shape parameter of gamma distribution | 0.3 | 0.3 | 0.3 | 0.3 |
| Dominance coefficient | 0.5 | 0.5 | 0.5 | 0.5 |

**Figure 1:**
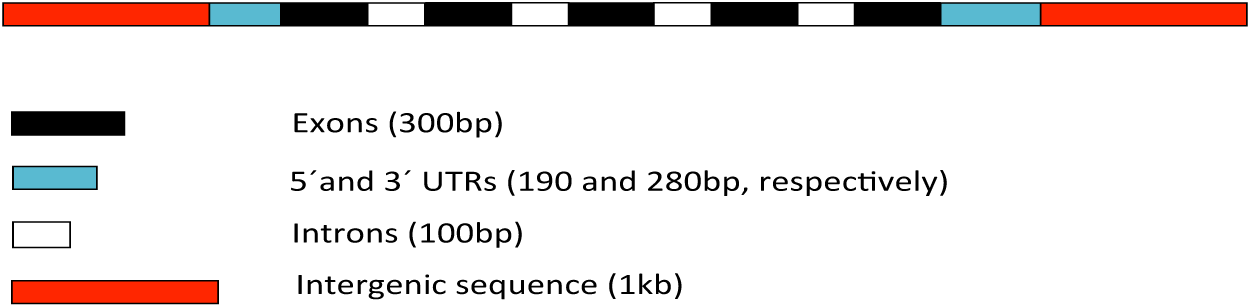
The gene model used in the simulations.

We chose an *N_e_*/*N* ratio of 532, equivalent to an *N_e_* of 1.33 million for the natural population. This value was based on the mean autosomal synonymous site diversity value of *π*= 0.018 for an African population and a mutation rate of *μ*= 4.5 x 10^-9^ (Campos *et al.* 2017), using the standard equilibrium formula *π= 4N_e_μ*for neutral variability under the infinite sites model (Kimura 1971), and assuming (rather conservatively) that mean diversity has been reduced by hitchhiking effects to 76% of its value in the absence of selection. The selection coefficients for favorable mutations, and the proportions of mutations that are favorable, were chosen to match mean values inferred from the relation between the synonymous diversity of a gene and its rate of protein sequence evolution by Campos *et al.* (2017). The details of the selection parameters used here are described in Table 1. Both favorable and deleterious mutations were assumed to be semidominant.

To model recombination, we mostly used 5 rates of reciprocal crossing over, which were multiples of the standard autosomal recombination rate in Drosophila, adjusted by a factor of ½ to take into account the absence of recombinational exchange in males (Campos *et al.* 2017). These ‘effective rates of crossing over’ span most of the observed range, and were 0.5 x 10^-8^, 1 x 10^-8^, 1.5 x 10^-8^, 2 x 10^-8^ and 2.5 x 10^-8^ cM/Mb, respectively, where 1 x 10^-8^ is the standard rate. We also ran simulations with no crossing over. The simulations were run with and without non-crossover associated gene conversion events, using a rate of initiation of conversion events of 1 x 10^-8^ cM/Mb for autosomes (after correcting for the lack of gene conversion in males) and a tract length of 440 bp. Given that SLiM models gene conversion by considering only the effects of conversion events that were initiated on one side of a given nucleotide site, this rate of initiation is one-half of the values estimated from the experiments of (Hilliker *et al.* 1994) and Miller *et al.* (2016), thus providing a conservative estimate of the effect of gene conversion. We did not vary the rate of initiation of gene conversion when using different rates of crossing over, since this rate appears to be fairly constant across the Drosophila genome, even in regions that lack crossing over (Langley *et al.* 2000; Comeron *et al.* 2012; Miller *et al.* 2016).

In addition to the simulations of autosomes, we ran simulations that were intended to represent X chromosomal mutations with equal fitness effects in the two sexes but with stronger selection than for autosomal mutations, as is expected on both theoretical and empirical grounds (Charlesworth *et al.* 2018). X-linked loci spend two-thirds of their time in females where they can recombine, so that the effective rates of crossing over and initiation of gene conversion events for X-linked loci should be 4/3 times the autosomal values for X-linked genes that have similar parameter values in females to the autosomal ones (Campos *et al.* 2013). The version of SLiM that we used did not permit explicit modeling of an X chromosome. We therefore used an autosomal model with a population size of 2500, but assumed that the true *N_e_*was three-quarters of that for the autosomes. Because *N* was kept constant, the autosomal rates of crossing over and initiation of gene conversion events were used in the simulations. In order to ensure that X-linked neutral variability in the absence of selection was three-quarters of the autosomal value, the mutation rate was multiplied by 3/4. Finally, with semi-dominance and equal fitness effects of mutations in males and females, the selection coefficient for an X-linked mutation is 4/3 times that for an autosomal mutation with the same selection coefficient, implying that the scaled selection coefficients are the same. To mimic stronger selection for positively selected mutations on the X chromosome, we therefore simply multiplied the scaled selection coefficients by a given factor, either 1.5 or 2. No adjustment was made to the scaled selection coefficient for deleterious mutations.

According to the number of genes simulated, we ran four sets of simulations with genomic regions of 20 (87.4 kb), 70 (305.9 kb), 140 (610 kb) and 210 (920 kb) genes. Most of our simulations used multiples of 70 genes because this represents a genomic region with a similar number of genes to the 4^th^ chromosome of *D. melanogaster*, which the simulations with zero crossing over are intended to model. Each simulation was run for 35000 (14*N*) generations, which is sufficient to allow the frequency distributions of neutral and deleterious mutations to reach equilibrium (see the online Supplementary Information, File S1, Figure S1). For the final estimates of diversity statistics (mean values of nucleotide site diversity, Tajima’s *D* and the proportions of singletons at synonymous, NS, intron and UTR sites) we used data from the final generation of each simulation. For calculating the numbers of fixations of favorable mutations at NS and UTR sites, we recorded the fixations that occurred during the last 20000 (8*N*) generations. In most cases, 20 replicate simulations were run for each parameter set, but a number of cases used 9 or 10 replicates.

Four different scenarios were simulated. First, purely neutral mutations were simulated in order to calculate the diversity statistics for the neutral reference. Three types of scenario with hitchhiking were simulated (i) SSWs only (ii) BGS only (iii) both SSWs and BGS. Sample sizes of 20 haploid genomes (a similar size to that used by Campos *et al.* 2017) were used for calculating the population genetic statistics. Mean values of each statistic over genes and replicate runs for a given model were recorded, with upper and lower 2.5 percentiles obtained by bootstrapping the mean values per gene of the chosen statistic across replicates (for brevity, we will refer to these as 95% confidence intervals). The statistics generated by the simulations are presented in the online Supplementary Information, Files S2 and S3.

No new data or reagents were generated by this research. Details of the mathematical derivations are described in the Supplementary Information, File S1. The detailed statistics for the results of the computer simulations are provided in the Supplementary Information, Files S2-S3. The code for the computer programs used in the models described below is available in the Supplementary Information, File S4.

## Theoretical Results

### Background selection

The predicted effect of BGS in a multi-site context can be described by the quantity *B* = exp(–*E*), where *B* is the ratio of expected neutral diversity at a focal neutral site under BGS to its value in the absence of BGS (which is equivalent to the corresponding ratio of mean coalescence times), and *E* is the sum of the effects of each selected site (Hudson and Kaplan 1995; Nordborg *et al.* 1996; Santiago and Caballero 1998). We assume a genomic region containing many genes, with selected sites that are continuously distributed with constant density, as in Model 3 of Charlesworth (2012b). We distinguish between nonsynonymous (NS) sites and untranslated regions regions (UTRs). This is, of course, a somewhat crude approximation, given that our genic model includes neutrally evolving intronic and intergenic sequences. For simplicity, we describe the case of autosomal inheritance, but parallel results hold for X-linked loci, with the appropriate changes in selection, mutation and recombination parameters.

We model both reciprocal exchange via crossing over and non-crossover associated gene conversion in the model. We assume that the main contribution from gene conversion comes from sites that are sufficiently distant that gene conversion causes recombination between them at a fixed rate *g* = *r_g_d_g_*(*r_g_*is the rate of initiation of gene conversion events in females and *d_g_*is the mean tract length). This is the limiting value of the general expression for the rate of recombination due to gene conversion for sites separated by *z* basepairs, *g*[1 – exp (– *z*/*d_g_*)] (Langley *et al.* 2000; Frisse *et al.* 2001), after correcting for the lack of gene conversion in male meiosis.

Because SLiM assumes no crossover interference, the relation between the frequency of crossing over and map distance in the simulations follows the Haldane mapping function (Haldane 1919), such that the frequency of crossing over between a pair of sites separated by *z* basepairs is given by:

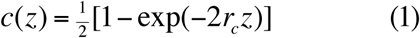

where *r_c_* is the rate of crossing over per basepair.

The net frequency of recombination between the sites is *r*(*z*) = *g* + *c*(*z*). The predicted value of *E* for a given selection coefficient, *t* = *hs*, against heterozygous carriers of a deleterious mutation, *E_t_*, is given by Equations S1 – S5 in section S1 of File S1. To obtain the final value of *E,* this equation is numerically integrated over the probability distribution of *t* values for NS and UTR sites separately, with total deleterious mutation rates *U_N_*and *U_U_*for NS and UTR sites, respectively, giving values *E_N_* and *E_U_* for the corresponding BGS effects.

To mimic the simulation results, we assume a gamma distribution with a shape parameter of 0.3. As in previous studies, we ignore all deleterious mutations with a scaled selection coefficient *γ* = 2*N*_e_s below a critical value *γ*_*c*_, in order to deal with the problem that very weakly selected mutations are subject to drift and contribute little to BGS effects (Nordborg *et al.* 1996). Following Nordborg *et al.* (1996) and Campos *et al.* (2017), we set *γ*_*c*_ = 5, and the gamma distributions for both NS and UTR mutations were truncated accordingly. Numerical results for the integral of the kernel of the gamma distribution from *γ*_*c*_to infinity allow the proportion of mutations that exceed *γ*_*c*_to be calculated; these are denoted by *P_N_*and *P_U_*for NS and UTR sites, respectively. With the parameters used in the simulations of autosomes, this gives *P_N_*= 0.871 and *P_U_*= 0.694. The final value for *E* is *P_N_E_N_*+ *P_U_E_U_*, from which *B* can be obtained as exp(– *E*).

### Selective sweeps

Various methods have been used to predict the approximate effect of a single selective sweep on diversity statistics at a partially linked neutral site in a randomly mating population, as well as for the associated distortion of the neutral site frequency spectrum at segregating sites (Maynard Smith and Haigh 1974; Kaplan *et al.* 1989; Stephan *et al.* 1992; Barton 1998, 2000; Gillespie 2000, 2001; Durrett and Schweinsberg 2004; Kim 2006; Pfaffelhuber *et al.* 2006; Coop and Ralph 2012; Bossert and Pfaffelhuber 2013). Here we present a simple heuristic derivation of the effect of a sweep on the pairwise neutral nucleotide site diversity, *π*, based on a combination of coalescent process and diffusion equation approaches. Following earlier approaches, we obtain the probability that a neutral lineage associated with a favorable allele at the end of a sweep was also associated with it at the start of the sweep, rather with the wild-type allele at the selected locus (Figure 2).

**Figure 2:**
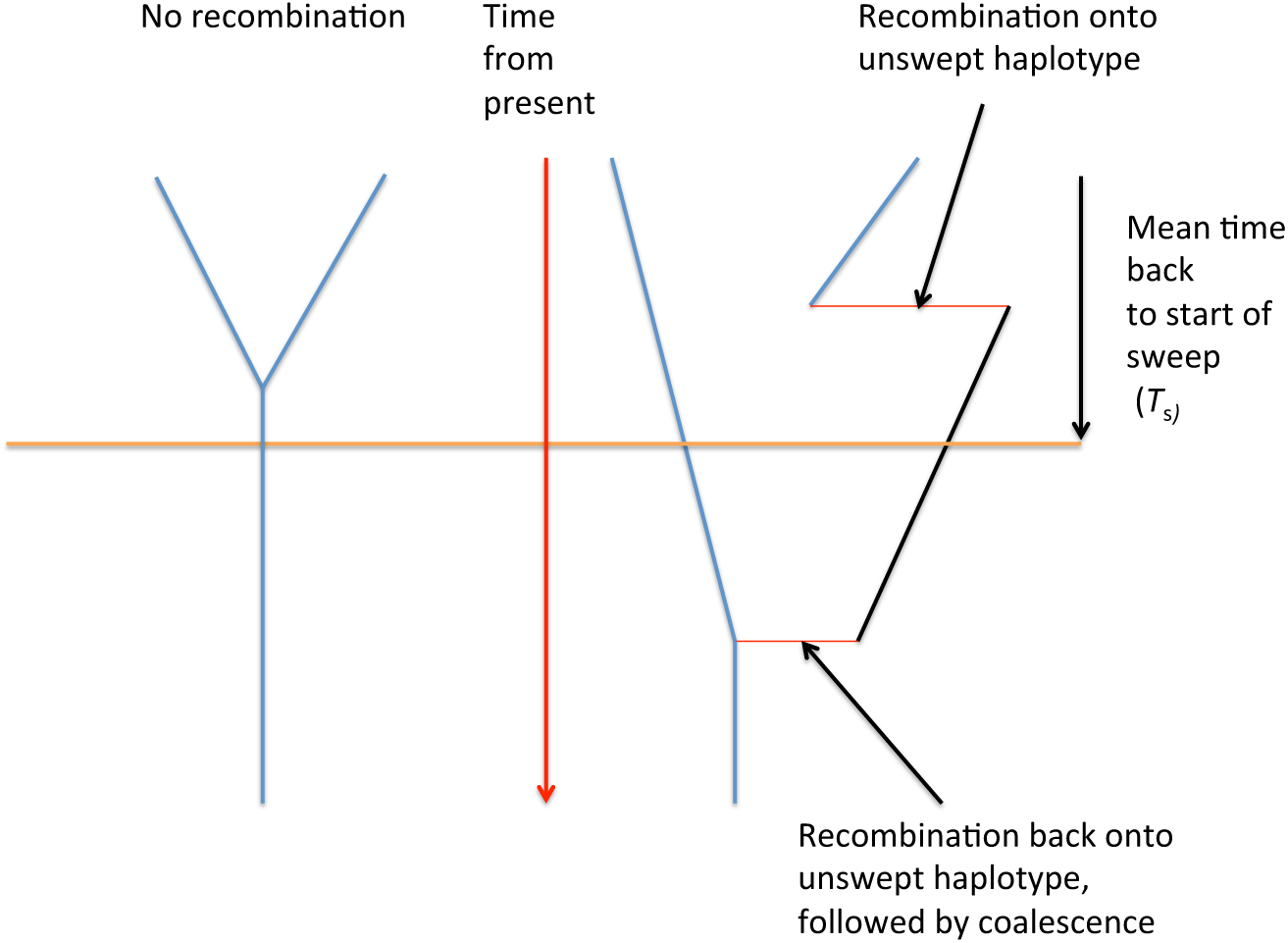
The possible fates of pairs of neutral lineages sampled after sweep, with no recombination on the left, and two recombination events on the left.

We consider separately the deterministic and stochastic phases of the spread of a favorable mutation, which were identified early in the history of the study of sweeps (Maynard Smith and Haigh 1974; Kaplan *et al.* 1989; Stephan *et al.* 1992; Barton 1998). The initial spread of a favorable allele A_2_from a frequency of 1/(2*N*) is subject to large stochastic effects. With semi-dominance, the probability that A_2_survives this effectively neutral period is approximately *Q* = *N_e_s/N* in a large population (Kimura 1962), assuming that the scaled selection coefficient, *γ* = 2*N_e_s*, is much greater than one (*s* is the selective advantage to homozygotes for the favorable mutation). As pointed out by Maynard Smith (1976), the overall expected frequency of A_2_during this quasi-neutral phase (including losses) is approximately 1/(2*N*), after which it starts to behave deterministically. The expected frequency of A_2_at the end of the quasi-neutral phase, conditioning on its surviving with probability *Q,* is thus 1/(2*NQ*) = *γ*^−1^. More rigorously, Martin and Lambert (2015) used branching process theory to show that the frequency of A_2_at the end of the first stochastic phase is exponentially distributed, with mean *γ*^−1^ and variance *γ*^−2^.

In the presence of BGS, we follow Kim and Stephan (2000) and assume that *N_e_* in the formula for fixation probability is multiplied by a constant, *B* (see above). As shown below, this constant is somewhat different for the effect of BGS on purely neutral processes, such as the level of neutral variability, and for the effect of BGS on the fixation of favorable mutations, since selected variants are more resistant to the effects of BGS than neutral variants (Johnson and Barton 2002). We denote these two constants by *B*_1_and *B*_2_, respectively, and write λ for the ratio *B*_1_/*B*_2_. The critical frequency at which A_2_can be treated as behaving deterministically is then (*B*_2_*γ*)^−1^, using the argument in the preceding paragraph. When A_2_reaches a frequency close to 1, there is a second stochastic phase in which it drifts to fixation fairly rapidly, as described below. We assume that all other effects of BGS are similar to those for neutral variability, with *B*_1_as the factor that multiplies *N_e_*.

The expectation of the time spent in the deterministic phase can be found as follows. As described by Ewens (2004, p.169), a semi-dominant favorable allele has the property that the expected time spent in a small interval of allele frequency *q* to *q*+ d*q* is the same as the time spent in the interval 1 – *q* to 1 – *q* – d*q*. This implies that the expected time that A_2_ spends between 1/(2*N*) and (*B_2_*γ**^−1^ is the same as the expected time it spends between 1 – (*B_2_*γ**)^−1^ and 1 – 1/(2*N*), so that *q* during the deterministic phase can conveniently be treated as lying between (*B_2_*γ**)^−1^ and 1 – (*B_2_*γ**)^−1^. Using the solution of the deterministic selection equation d*q*/d*t* = ½ *spq* for a semi-dominant allele Haldane (1924), the expected time spent in this interval (expressed in units of coalescent time, 2*N_e_*generations) ≈ 2*γ*^−1^ ln(*B*_2_^2^*γ*^2^) = 4*γ*^−1^ ln(*B_2_*γ**).

The expected times spent in the two stochastic phases can be found as follows. Using Equation 16 of Kimura and Ohta (1973) and the fact that *N_e_*is multiplied by *B*_1_to take BGS into account, the expected first passage time of a neutral allele from initial frequency 1/(2*N*) to a frequency *q* is:

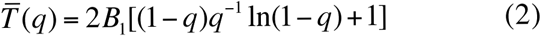

For *q* << 1, this time is approximately equal to *B*_1_*q*, so that the additional expected time spent in the first stochastic phase is approximately λ*γ*^−1^. By the above symmetry argument, the same applies to the time between 1 – (*B_2_*γ**^−1^ and 1 – 1/(2*N*). The total expected time to fixation of A_2_when *γ* >> 1 is thus:

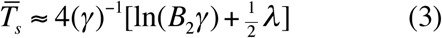

This expression is very close to Equation A17 of Hermisson and Pennings (2005) for the case with *B*_1_= *B*_2_= 1, which was derived directly from the diffusion equation for the mean sojourn time of a favorable mutation in a finite population.

As far as the effect of a substitution on neutral diversity is concerned, we note that the rate (in units of coalescent time) at which a neutral lineage that is associated with A_2_at time *T* recombines onto a background of A_1_is *p*(*T*)*ρ*, where *p*(*T*) is the frequency of the wild-type allele at time *T* and *ρ* = 2*N_e_r* is the scaled recombination rate. Here, *T* = 0 at the time of fixation of the favorable allele, and *T = T_s_*at the time when it arose in the population. From the symmetry of the selection equation, the mean frequency of A_1_over the deterministic phase is 0.5, so that that *ρ* should be discounted by a factor of ½ during this part of the process. (Note that this argument ignores the possibility of coalescence competing with recombination during the sweep, as pointed out to us by Matthew Hartfield. A more rigorous treatment that includes such competition will be presented elsewhere.)

For sample paths in which A_2_reaches the critical frequency (*B_2_*γ**^−1^, the expected duration of the first stochastic phase is equal to the expected value of the first passage time to this frequency, *λ*γ**^−1^, and its variance is *λ*^2^*γ*^−2^/3 (File S1, section S2). During this period, a single lineage recombines with A_1_haplotypes at a rate close to *ρ*, since A_1_dominates the population, thus contributing 2*ρ* (*λ*γ**^−1^ + *δT_s_*_1_) to the mean number of recombination events, where *δT_s_*_1_is the departure of the duration of the first stochastic phase from its expectation. The final stochastic phase has effectively zero probability of contributing to recombination, due to the prevalence of the favored allele, and can be ignored for this purpose.

We exploit the fact that the frequency of A_2_at the end of the first stochastic phase is exponentially distributed (Martin and Lambert 2015) to show that the variance of the time to fixation caused by fluctuations in the initial frequency of A_2_at the start of the deterministic phase yields an additional variance term of 16(*γ*)^−2^ (File S1, section S2). Since this phase has a mean frequency of A_2_of 0.5, the relevant product of the recombination rate and a fluctuation in deterministic sweep time (*δT_s_*_2_) is *ρδT_s_*_2_rather than 2*ρ δT_s_*_2_.

The probability *P_cs_*that the two sampled haplotypes coalesce as a result of the sweep is equivalent to the probability that neither member of a pair of haplotypes sampled at time *T* = 0 recombined onto an A_1_background, provided that the sweep durations are so short that no coalescence can occur among non-recombined haplotypes during the sweep (Wiehe and Stephan 1993). This probability is given by the first term of a Poisson distribution, whose mean is equal to the expected number of recombination events over the duration of a substitution. We thus have:

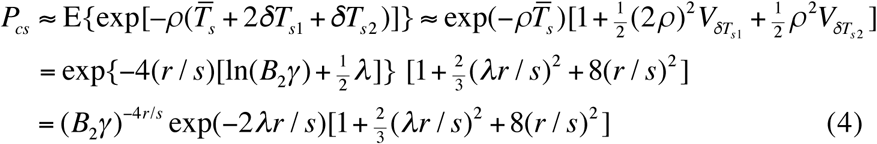

The term (*B*_2_*γ*)^−4*r*/*s*^in the third line of Equation 4 is the deterministic phase contribution to the effect of a sweep, first derived by Barton (2000, 1998) for the case of *B*_1_= *B*_2_= 1, using a more rigorous approach. It has been used in several subsequent studies (Weissman and Barton 2012; Elyashiv *et al.* 2016; Campos *et al.* 2017). The last term is second-order in *r*/*s* and is likely to be of minor importance, since sweeps only have substantial effects on variability when *r/s* << 1. The second term has a somewhat larger effect; e.g. with no BGS, *γ* =100, and *r/s* = 0.1, it reduces *P_cs_* from 0.158 to 0.130. An extension to these results is described in section S3 of File S1 (Equation S20), which allows for multiple recombination events that bring a recombined lineage back onto an A_2_background (Figure 2).

### Sweeps at multiple sites

We now consider the effects of recurrent sweeps at multiple sites. The standard approach assumes that substitutions of favorable alleles are sufficiently rare that their effects on a given site can be treated as mutually exclusive events (Kaplan *et al.* 1989; Wiehe and Stephan 1993; Kim and Stephan 2000; Kim 2006), and this assumption will also be made here. We consider only a single gene, which is reasonable for favorable mutations whose selection coefficients are less than the rate of recombination between sites in different genes, as is usually the case here. Both these assumptions are supported by the the simulation results presented below, except for cases with very low rates of recombination.

We use the expression for the probability of a sweep-induced coalescent derived above (Equation 4) to obtain an approximate expression for the net rate of coalescent events experienced at a given neutral site (in units of 2*N_e_* generations), due to recurrent selective sweeps at NS and UTR sites:

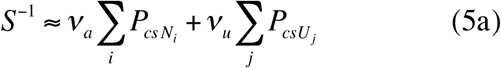

where *ν_a_* and *ν_u_* are the rates (in units of coalescent time) at which substitutions of favorable mutations occur at NS and UTR sites respectively; *P_cs Ni_*and *P_cs Uj_*are the rates of sweep-induced coalescent events induced by the *i*th NS site and *j*th UTR site, respectively. The summations are taken over all the sites in the gene that are under selection. The notation *S*^−1^ is used to denote the reciprocal of the expected time to coalescence due to sweeps, *S*.

Using Equations 4 and S20, we have:

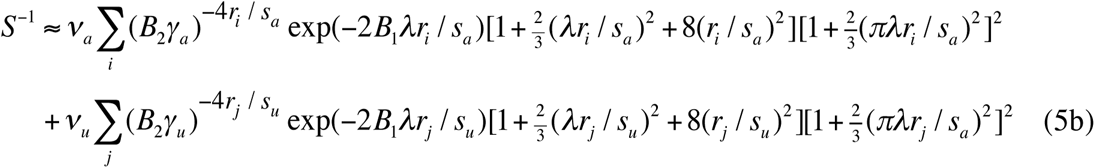

where subscripts *a* and *u* denote NS and UTR mutations, respectively.

If we assume that the fixation probability of a favorable mutation in the presence of BGS is discounted by a factor of *B*_2_compared with the standard value (see above), we have:

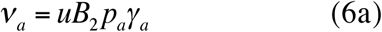

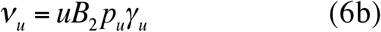

where *u* is the mutation rate per nucleotide site, and *p_a_*and *p_u_*are the proportions of all new NS and UTR mutations, respectively, that are selectively favored.

The summation formula used in the sweep calculations assumes that every third basepair in an exon is a neutral site, with the other two being subject to selection (Campos *et al.* 2017). This differs from the SLiM procedure of randomly assigning selection status to exonic sites, with a probability *p_s_* of being under selection (*p_s_* = 0.7 in the simulations used here). To correct for this, the overall rate of NS substitutions in Equations 5 was adjusted by multiplying by 0.7 x 1.5. Since we are confining ourselves to a single gene, it is reasonable to assume a linear genetic map. The crossing over contribution to *r_i_*is then given by *r_c_z_i_*, where *z_i_*is the physical distance between the neutral and selected sites. There is also a contribution from gene conversion, as described in the section on modeling BGS.

Following Kaplan *et al.* (1989), Wiehe and Stephan (1993) and Kim and Stephan (2000), coalescent events caused by selective sweeps and coalescent events caused by neutral drift can be considered as competing exponential processes with rates *S*^−1^ and *B*_1_^−1^, respectively, on the coalescent timescale of 2*N_e_* generations. Under the infinite sites model (Kimura 1971), the ratio of expected nucleotide site diversities at a neutral site, relative to the value in the absence of selection at linked sites (θ = 4*N_e_u* where *u* is the neutral mutation rate per basepair), can then be written as the expected time to coalescence, when time is measured in units of 2*N_e_* generations:

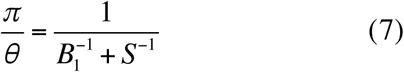

The simulation results for synonymous site nucleotide site diversities were presented as mean values over all genes in the region simulated. Since we are modeling only a single gene, the mean of *π*/θ in Equation 7 over all synonymous sites in a gene should be used for comparison with the simulation results. In practice, the values obtained by substituting the mean value of *S*^−1^ across synonymous sites into Equation 7 give almost identical results, and this is used for results described below.

### The effect of sweep duration on mean coalescent time

Equation 7 assumes that the duration of a sweep is negligible in comparison to the times between successive sweeps and to the mean neutral coalescent time 2*N_e_*, so that sweeps can be treated as point events. This assumption is, however, violated if selection is sufficiently weak. For example, with *γ* = 250 the deterministic component of the duration of an adaptive substitution given by Equation 3 is approximately 10% of the coalescent time. The assumption that the entire time between sweep-induced coalescent events is available for neutral coalescent events is therefore only an approximation, and leads to an underestimate of the effects of sweeps when these are frequent.

Here we develop an alternative approach that uses the mean diversity between successive substitutions as an estimate of the expected diversity under recurrent sweeps. This is likely to overestimate the effects of sweeps compared with the mean for randomly sampled time points, but the simulation results described below show that the resulting expressions (Equations 12) provide a good fit. We assume that adaptive substitutions occur in a gene at a constant rate *ω* per unit of coalescent time, given by the sum over the rates per site for the NS and UTR sites in the gene. This quantity can be found from Equations 6 by multiplying ν_*a*_ by 70% of the number of NS sites in a gene, and ν_*u*_ by the number of UTR sites. We then look back in time, and evaluate the time average of the divergence of *π/θ* from its equilibrium value over the period since the previous substitution.

To do this, we denote the expected neutral diversity at a neutral site immediately after a substitution by *π*_0_, and the expected neutral diversity at the time of initiation of a new substitution by *π*_1_. We have:

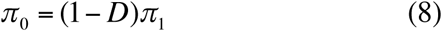

where *D* is the probability that each member of a pair of lineages carrying the favorable mutation has failed to recombine during the substitution, conditioned on the completion of a substitution. Because the expected reduction in neutral diversity due to recurrent sweeps is *S*^−1^, given by Equation 5b, we have *D* = (*ωS*) ^−1^, thereby establishing the relationship between *π*_0_and *π*_1_(the assumption that the coalescent time for the pair of swept lineages is zero is relaxed below).

Under the infinite sites model (*θ* << 1), the equilibrium diversity in the absence of sweeps is *B*_1_*θ*. In this case, the standard formula for the rate of approach of neutral diversity to its equilibrium value (Malécot 1969, p.40; Wiehe and Stephan 1993, Equation 6a), gives the following expression for the diversity at a time *T* after a substitution:

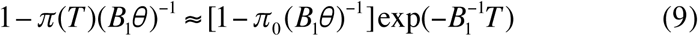

(The factor of *B*_1_^−1^ in the exponent reflects the reduction in *N_e_*caused by BGS, resulting in a corresponding acceleration in the rate of approach to equilibrium.)

The expected diversity over the relevant period, *π*, is thus given by:

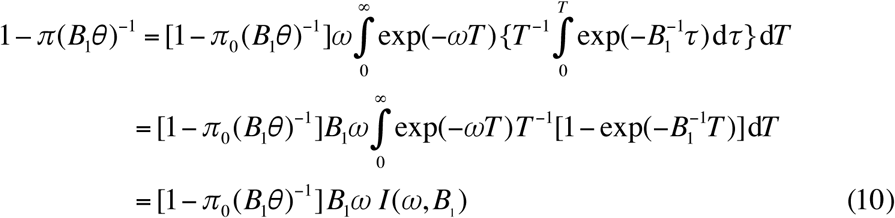

Formulae for *I*(*ω*, *B*_1_) are derived in File S1, section 5.

Furthermore, *π*_1_is given by:

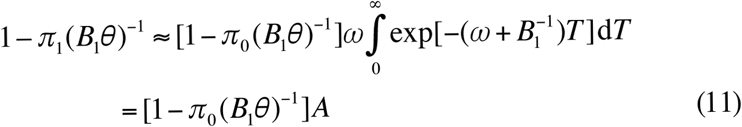

where *A* =*ω*/(*ω* + *B*_1_^−1^).

In the absence of any recovery of diversity during the sweep itself, Equations 8-10 together yield the final expression:

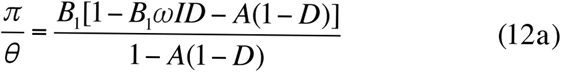

In the limit as *ω* approaches zero, *ωI* and *A* both tend to 0, and *AD* tends to *B*_1_*S*^−1^. The value of *π/θ* for small *ω* is thus approximately 1/(*B*_1_+ *S*^−1^), corresponding to Equation 7.

To allow for a non-zero mean time to coalescence during the sweep, *T_cs_*, the post-sweep diversity *π*_0_is modified by adding *DT_cs_θ* to Equation 8, where *T_cs_*is given by Equation S10 (this is an underestimate, since it ignores recombination during the sweep). This adds a small additional component to Equation 12a, giving:

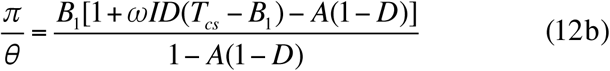

Equations (12) assume that the sample is taken in an interval between two successful sweeps. A correction can be applied to take into account the possibility that a sample is taken during a sweep; this effect is expected to be small unless sweep-induced coalescents are very frequent and the time occupied by a sweep is relatively large compared with the neutral coalesencent time (File S1, section S6).

### Continuum approximation for effects of recurrent sweeps

A useful approximation can be obtained by treating a gene as a continuum, following the treatment of BGS in Campos *et al.* (2017) and selective sweeps by Weissman and Barton (2012). We correct for the effect of introns simply by reducing the density of NS sites in the coding sequence. This is done by multiplying the density within exons by the fraction of the sites that are exons among the total length of exons, introns and UTRs. In addition, we approximate the effect of gene conversion by writing the net recombination rate between sites separated by *z* basepairs as (*r_c_* + *g_c_*)*z* when *z ≤ d_g_*, and as *r_c_ z* + *g* (where *g* = *g_c_d_g_*) when *z > d_g_* (Andolfatto and Nordborg 1998). The resulting expressions for sweep effects are derived in File S1, section S7. These do not include any corrections for multiple recombination events or for the variances in the first stochastic phase and deterministic phase durations, since these make the integrations analytically intractable.

## Simulation Results

### Effects of background selection alone

Table 2 shows simulation results with BGS alone, using the gene model described in the Material and Methods, for chromosomal regions with varying numbers of autosomal loci and rates of crossing over, with and without gene conversion at the standard rate. The estimates of *B*_1_, the ratio of the mean synonymous site nucleotide diversity to the value without selection (θ), are shown in the table, together with their 95% confidence intervals (CIs) over replicate simulations. The mean value of θ from simulations of neutral mutations in the absence of selection at linked sites was 0.0223, with 95% CI (0.0227, 0.0229), which is slightly lower than the theoretical value on the infinite sites model (0.0239), presumably due to the slight deviations from the infinite sites assumption in SLiM. The ratios of the simulated synonymous site diversities to 0.0223 were used for the estimates of *B*_1_ in the table.. Table S1 of File S1 shows comparable results for the model of X-linked loci summarized in Table 1, assuming intermediate dominance and a mean scaled selection coefficient against homozygous deleterious mutations and shape parameter that are equal to the autosomal values.

**Table 2:** BGS predictions and simulation results for autosomal values of *B*_1_ = *π/θ*.

| Xover Rate | 0.5 | 1.0 | 1.5 | 2.0 | 2.5 |
| --- | --- | --- | --- | --- | --- |
| **No g.c** |  |  |  |  |  |
| 20 genes | 0.687, 0.723<br>(0.702, 0.745) | 0.798, 0.838<br>(0.823, 0.852) | 0.845, 0.861<br>(0.843, 0.879) | 0.871, 0.868<br>(0.849, 0.887) | 0.888, 0.913<br>(0.894, 0.933) |
| 70 genes | 0.593, 0.643<br>(0.623, 0.645) | 0.716, 0.737<br>(0.721, 0.750) | 0.767, 0.790<br>(0.770, 0.799) | 0.794, 0.818<br>(0.810, 0.827) | 0.812, 0.830<br>(0.824, 0.838) |
| 140 genes | 0.514, 0.543<br>(0.534, 0.550) | 0.632, 0.655<br>(0.648, 0.663) | 0.679, 0.701<br>(0.694, 0.709) | 0.704, 0.723<br>(0.717, 0.729) | 0.719, 0.731<br>(0.724, 0.739) |
| 210 genes | 0.452, 0.489<br>(0.481, 0.491) | 0.559, 0.582<br>(0.574, 0.590) | 0.601, 0.620<br>(0.618, 0.624) | 0.623, 0.642<br>(0.638, 0.646) | 0.637, 0.654<br>(0.650, 0.658) |
| **G.c.** |  |  |  |  |  |
| 20 genes | 0.753, 0.796<br>(0.775, 0.819) | 0.836, 0.883<br>(0.862, 0.903) | 0.872, 0.905<br>(0.889, 0.920) | 0.892, 0.907<br>(0.887, 0.929) | 0.905, 0.924<br>(0.905, 0.942) |
| 70 genes | 0.650, 0.686<br>(0.677, 0.693) | 0.750, 0.782<br>(0.767, 0.799) | 0.791, 0.816<br>(0.797, 0.834) | 0.813, 0.820<br>(0.813, 0.827) | 0.827, 0.838<br>(0.830, 0.848) |
| 140 genes | 0.563, 0.594<br>(0.588, 0.601) | 0.662, 0.687<br>(0.676, 0.696) | 0.700, 0.719<br>(0.707, 0.728) | 0.720, 0.725<br>(0.720, 0.731) | 0.733, 0.736<br>(0.729, 0.744) |
| 210 genes | 0.496, 0.525<br>(0.519, 0.531) | 0.586, 0.605<br>(0.598, 0.613) | 0.620, 0.640<br>(0.635, 0.645) | 0.638, 0.639<br>(0.635, 0.640) | 0.649, 0.657<br>(0.652, 0.661) |

The left-hand upper entries in the cells show the predicted values of $B_1$ , the ratio of the mean synonymous site diversity with BGS (but no sweeps) to its value in the absence of BGS (using Equations S1, S2, S3, and S5, integrated over the truncated gamma distribution).
The right-hand upper entries are the corresponding observed mean values.
The lower entries are the lower and upper 2.5 percentiles of the observed values of $B_1$ , obtained from the means of the synonymous site diversities over the entire region for each replicate simulation.
The rows labelled 'Xover rate' refer to the results for rates of crossing over with ratios of 0.5, 1, 1.5, etc. to the standard rate of $5.32 \times 10^{-6}$ used in the simulations.
Cases with no gene conversion are denoted by 'No g.c.' and cases with the standard gene conversion parameters by 'G.c.'.

Tables 2 and S1 also show the predicted values of *B*_1_ using the continuum model of BGS with the Haldane mapping function described above, using the formulae in File S1, section S1. To obtain these values, Equation S3 was numerically integrated over the gamma distribution of selection coefficients, truncated at *γ*_*c*_= 5 (see the Material and Methods). The theoretical predictions for the X-linked case are equivalent to those for a mutation rate of ¾ times the autosomal values, with the same values as the autosomal case for all other parameters. Overall, there is a fairly good fit between the theoretical predictions and the simulation results, although the theoretical values of *B*_1_ are mostly slightly smaller than the simulation values, probably because intergenic sequences have been ignored.

However, if the additional term in *E* contributed from neutral mutations that arise in repulsion from a linked deleterious mutation (Equations S1b, S5d and S5e) is ignored, the fits are much less good, especially for the higher rates of crossing over and larger numbers of genes. For example, with 70 autosomal genes and the standard rate of gene conversion, the predicted values of *B*_1_ are then 0.681, 0.790, 0.835, 0.860 and 0.875 for crossover rate factors of 0.5, 1, 1.5, 2 and 2.5, respectively. With 210 genes, the corresponding *B*_1_ values are 0.583, 0.696, 0.739, 0.762 and 0.776; the last value is 20% larger than when the additional term is included.

Similarly, use of a linear relation between physical distance and map distance, which has been assumed in most theoretical models of BGS, generally gives a poorer fit to the results for the higher rates of crossing over (Table S2 of File S1), except when the number of genes and the map length of the region are both small, reflecting the effect of double crossing over in reducing the net rate of recombination between distant sites. Nonetheless, the fit is surprisingly good overall; indeed, the linear map predictions using Equations S2c, S2d, S4, S5d and S5e often provide a better fit to the simulation results for the cases with 20 and 70 genes. The implications of these effects of the inclusion of the repulsion mutations, and the difference between the linear and Haldane maps, are considered in the Discussion.

### Effects of background selection on the rate of fixation of favorable mutations

The main goal of our work is to analyse the joint effects on neutral diversity of BGS and SSWs, and the extent to which these can be predicted by the relatively simple Equations 7 and 12. A core assumption behind these equations is that the fixation probability of a new favorable mutation is affected by BGS as though *N_e_* is multiplied by a factor that is equal or close to the value that applies to neutral diversity (Kim and Stephan 2000).

We have tested this assumption by comparing the mean numbers of fixations of favorable mutations observed over the last 15,000 (8*N*) generations of the simulations, both without BGS and with BGS. The ratio of these means provides a measure of *B* (*B*_2_) that can be compared to the value of *B* estimated from neutral diversity (*B*_1_). There are two reasons why we would not expect perfect agreement. First, a sufficiently strongly selected favorable variant could resist elimination due to its association with deleterious mutations, and instead might drag one or more of them to high frequencies or fixation (Johnson and Barton 2002; Hartfield and Otto 2011). Second, the incursion of selectively favorable mutations may perturb linked deleterious mutations away from their equilibrium, even if they do not cause their fixation.

Such Hill-Robertson interference effects (Hill and Robertson 1966; Felsenstein 1974) reduce the *N_e_*experienced by deleterious mutations, and hence their nucleotide site diversity, which is correlated with the mean number of segregating deleterious mutations. This reduction in the number of segregating deleterious mutations reduces the effects of BGS on incoming favorable mutations. For both these reasons, *B*_1_ is likely to be smaller than *B*_2_. Table S3 of File S1 provides evidence that the mean number of segregating deleterious mutations is indeed reduced by selective sweeps, except for the cases with no crossing over, for which the rate of sweeps is greatly reduced compared with cases with crossing over.

The results for autosomal loci in Table 3 show that BGS has a substantial effect on the rate of adaptive substitutions (Table S4 of File S1 presents some parallel results for X-linked loci). The most extreme case is when there is no crossing over, a regime in which the efficacy of BGS is undermined by Hill-Robertson interference among the deleterious mutations, so that the assumptions underlying the BGS equations tested in the previous section are violated (McVean and Charlesworth 2000; Comeron and Kreitman 2002; Kaiser and Charlesworth 2009; Seger *et al.* 2010; Good *et al.* 2014; Hough *et al.* 2017). For example, *B*_1_ for 70 genes with gene conversion is 0.086, close to the value found by Kaiser and Charlesworth (2009) for a similar sized region, whereas the standard BGS prediction is 0.0004. In contrast, the *B*_2_ values for favorable NS and UTR mutations are 0.26 and 0.28, respectively, approximately three times greater. This still represents a massive reduction in the efficacy of selection on favorable mutations, consistent with the evidence that their rates of substitution in non-crossover regions of the Drosophila genome are much lower than elsewhere (Charlesworth and Campos 2014).

**Table 3:**
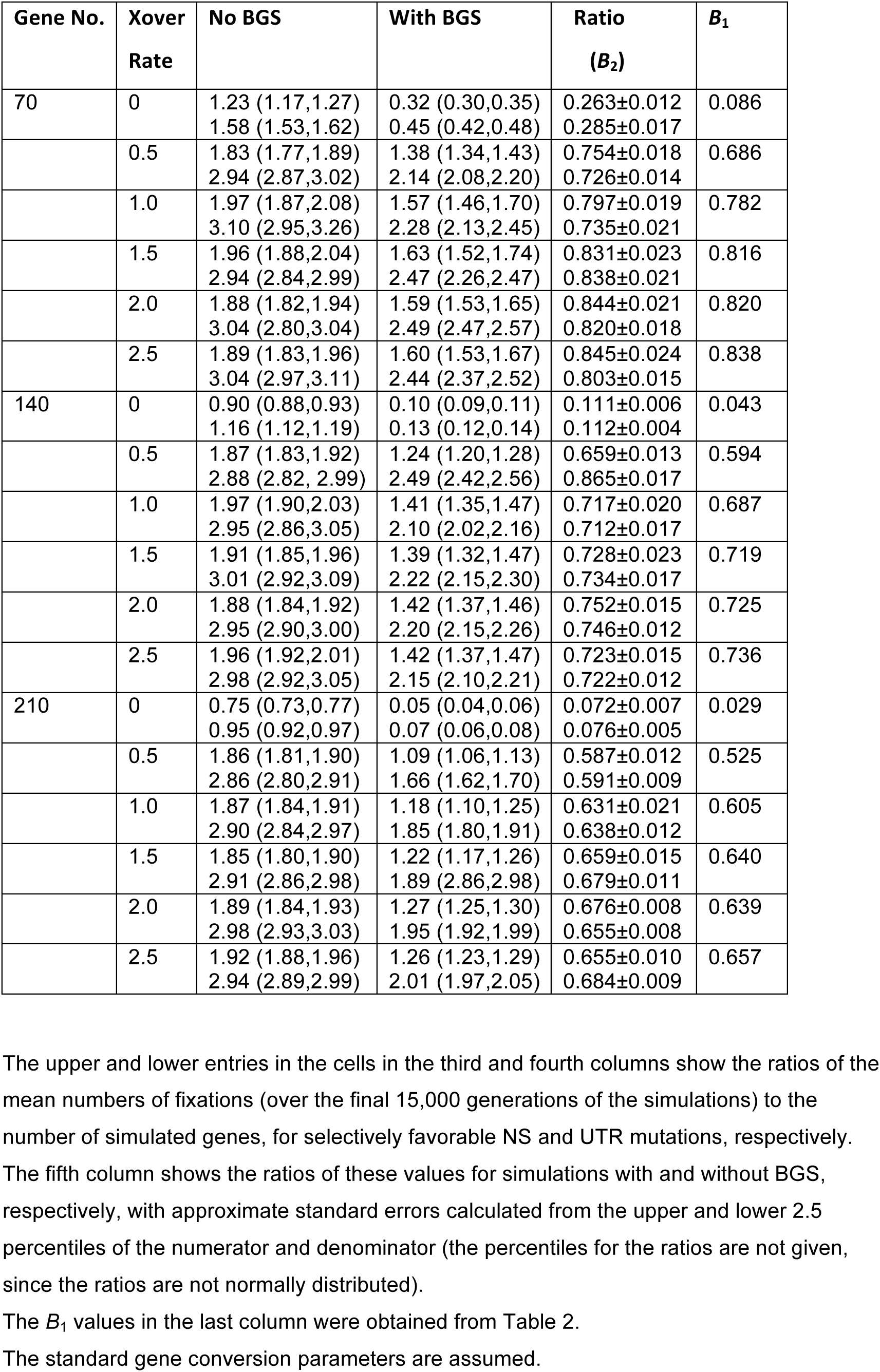
The effect of BGS on the numbers of fixations of selectively favorable autosomal mutations.

| Gene No. | Xover Rate | No BGS | With BGS | Ratio ($B_2$) | $B_1$ |
| --- | --- | --- | --- | --- | --- |
| 70 | 0 | 1.23 (1.17,1.27) | 0.32 (0.30,0.35) | 0.263±0.012 | 0.086 |
|  |  | 1.58 (1.53,1.62) | 0.45 (0.42,0.48) | 0.285±0.017 |  |
|  | 0.5 | 1.83 (1.77,1.89) | 1.38 (1.34,1.43) | 0.754±0.018 | 0.686 |
|  |  | 2.94 (2.87,3.02) | 2.14 (2.08,2.20) | 0.726±0.014 |  |
|  | 1.0 | 1.97 (1.87,2.08) | 1.57 (1.46,1.70) | 0.797±0.019 | 0.782 |
|  |  | 3.10 (2.95,3.26) | 2.28 (2.13,2.45) | 0.735±0.021 |  |
|  | 1.5 | 1.96 (1.88,2.04) | 1.63 (1.52,1.74) | 0.831±0.023 | 0.816 |
|  |  | 2.94 (2.84,2.99) | 2.47 (2.26,2.47) | 0.838±0.021 |  |
|  | 2.0 | 1.88 (1.82,1.94) | 1.59 (1.53,1.65) | 0.844±0.021 | 0.820 |
|  |  | 3.04 (2.80,3.04) | 2.49 (2.47,2.57) | 0.820±0.018 |  |
|  | 2.5 | 1.89 (1.83,1.96) | 1.60 (1.53,1.67) | 0.845±0.024 | 0.838 |
|  |  | 3.04 (2.97,3.11) | 2.44 (2.37,2.52) | 0.803±0.015 |  |
| 140 | 0 | 0.90 (0.88,0.93) | 0.10 (0.09,0.11) | 0.111±0.006 | 0.043 |
|  |  | 1.16 (1.12,1.19) | 0.13 (0.12,0.14) | 0.112±0.004 |  |
|  | 0.5 | 1.87 (1.83,1.92) | 1.24 (1.20,1.28) | 0.659±0.013 | 0.594 |
|  |  | 2.88 (2.82, 2.99) | 2.49 (2.42,2.56) | 0.865±0.017 |  |
|  | 1.0 | 1.97 (1.90,2.03) | 1.41 (1.35,1.47) | 0.717±0.020 | 0.687 |
|  |  | 2.95 (2.86,3.05) | 2.10 (2.02,2.16) | 0.712±0.017 |  |
|  | 1.5 | 1.91 (1.85,1.96) | 1.39 (1.32,1.47) | 0.728±0.023 | 0.719 |
|  |  | 3.01 (2.92,3.09) | 2.22 (2.15,2.30) | 0.734±0.017 |  |
|  | 2.0 | 1.88 (1.84,1.92) | 1.42 (1.37,1.46) | 0.752±0.015 | 0.725 |
|  |  | 2.95 (2.90,3.00) | 2.20 (2.15,2.26) | 0.746±0.012 |  |
|  | 2.5 | 1.96 (1.92,2.01) | 1.42 (1.37,1.47) | 0.723±0.015 | 0.736 |
|  |  | 2.98 (2.92,3.05) | 2.15 (2.10,2.21) | 0.722±0.012 |  |
| 210 | 0 | 0.75 (0.73,0.77) | 0.05 (0.04,0.06) | 0.072±0.007 | 0.029 |
|  |  | 0.95 (0.92,0.97) | 0.07 (0.06,0.08) | 0.076±0.005 |  |
|  | 0.5 | 1.86 (1.81,1.90) | 1.09 (1.06,1.13) | 0.587±0.012 | 0.525 |
|  |  | 2.86 (2.80,2.91) | 1.66 (1.62,1.70) | 0.591±0.009 |  |
|  | 1.0 | 1.87 (1.84,1.91) | 1.18 (1.10,1.25) | 0.631±0.021 | 0.605 |
|  |  | 2.90 (2.84,2.97) | 1.85 (1.80,1.91) | 0.638±0.012 |  |
|  | 1.5 | 1.85 (1.80,1.90) | 1.22 (1.17,1.26) | 0.659±0.015 | 0.640 |
|  |  | 2.91 (2.86,2.98) | 1.89 (2.86,2.98) | 0.679±0.011 |  |
|  | 2.0 | 1.89 (1.84,1.93) | 1.27 (1.25,1.30) | 0.676±0.008 | 0.639 |
|  |  | 2.98 (2.93,3.03) | 1.95 (1.92,1.99) | 0.655±0.008 |  |
|  | 2.5 | 1.92 (1.88,1.96) | 1.26 (1.23,1.29) | 0.655±0.010 | 0.657 |
|  |  | 2.94 (2.89,2.99) | 2.01 (1.97,2.05) | 0.684±0.009 |  |

The upper and lower entries in the cells in the third and fourth columns show the ratios of the mean numbers of fixations (over the final 15,000 generations of the simulations) to the number of simulated genes, for selectively favorable NS and UTR mutations, respectively.
The fifth column shows the ratios of these values for simulations with and without BGS, respectively, with approximate standard errors calculated from the upper and lower 2.5 percentiles of the numerator and denominator (the percentiles for the ratios are not given, since the ratios are not normally distributed).
The $B_1$ values in the last column were obtained from Table 2.
The standard gene conversion parameters are assumed.

For the other rates of crossing over, there is much closer agreement between the two estimates of *B*, although we always have *B*_1_ > *B*_2_. The discrepancy is largest for crossover rates of one-half the standard value, and seems to level off after the standard rate. As might be expected, it is smaller in the presence of gene conversion.

### Effects of interference among favorable mutations on their rates of substitution

With no recombination, Hill-Robertson interference among adaptive subsititutions is likely to be important, and makes analytical models of substitution rates much harder to develop. The effects of such interference can be predicted using the approximate Equation 4 of Neher (2013), based on Equation 39 of Desai and Fisher (2007). When this is adapted for the case of diploids with semidominance with *s* >> *U_b_*, the rate of substitution of favorable mutations, *ω*, is equal to 0.5*s* ln(*Ns*)/[ln(2*U_b_*/*s*)]^2^, where *s* is the homozygous selection coefficient for a favorable mutation, and *U_b_* is the net mutation rate to favorable mutations for the region. Combining NS and UTR mutations (these have similar selection coefficients in our simulations) and putting *s* = 0.05, *U_b_* = 0.00436 and *N* = 2500, *ω* = 0.00406; the ratio of *ω* to the baseline substitution rate in the absence of interfererence is 0.163.

The observed ratio of the rates of substitution for relative rates of crossing over of 0 and 2.5, with 70 genes and no gene conversion and no BGS was equal to 0.235, suggesting that the effect of interference is overpredicted by the approximation. Gene conversion increases the ratio to 0.570 (Table 3), so that it greatly reduces interference when crossing over is absent. BGS thus seems to play a more important role than SSWs in reducing the rate of substitution of favorable mutations when crossing over is absent, especially in the presence of gene conversion, as suggested by Campos *et al.* (2014). The properties of genomic regions with very low rates of crossing over will be analysed in more detail in a later publication.

In the absence of BGS, but with non-zero rates of crossing over, Tables 3 and S4 show little effect of the crossing over rate on the rate of fixation of favorable mutations. At first sight, this suggests that there is little interference among selectively favorable mutations with a rate of crossing over of one-half or more of the the standard rate. However, there is indirect evidence for such interference effects, from estimates of the extent of underdispersion of the numbers of adaptive substitutions observed over the last 8*N* generations of the simulations compared with the expectation for a Poisson distribution, as described in File S1, section S8. Here, underdispersion is measured by the ratio of the variance to the mean of the number of substitutions over the period of observation (Sellers and Morris 2017).

This analysis shows that interference causes a small loss of substitutions, leading to a reduction in the extent of the reduction in diversity caused by selective sweeps for the cases with crossing over, with approximately 5.5% of substitutions being lost due to interference. An approximate correction for interference can be made by multiplying the substitution rates for both NS and UTR mutations by the estimated proportion of substitutions that survive interference, although this ignores some of the complexities associated with the effects of interference on diversity (Kim and Stephan 2003; Chevin *et al.* 2008). In addition, it should be noted that the existence of underdispersion implies that the Poisson model of sweeps that is usually assumed is not exact, as pointed out by Gillespie (2001), introducing a further source of error into the predictions.

### Effects of selective sweeps on neutral diversity

This section is concerned with four main questions. First, to what extent does treating sweeps as point events affect the predictions of models of recurrent sweeps? Second, how well does the integral approximation for SSWs perform (Equations S24-S33), compared with the more exact summation formulae (Equations 5 and 6). Third, how well do the competing coalescent process approximations for the joint effects of BGS and SSWs perform, when the various corrections described above have been included? Finally, is less accuracy obtained by using the neutral BGS value (*B*_1_) instead of *B*_2_in the formulae for the effect of BGS on the fixation probability of a favorable mutation?

Table 4 presents the results of simulations with 70 autosomal genes, together with the predictions for the integral and summation formulae, with and without the corrections (the correction for interference described above is applied to all these cases). In the case of the corrected summation formulae, all the corrections described above were applied; for the integral results, only the corrections for expected sweep duration and interference were used. Parallel results for X-linked genes are shown in Table S6 of File S1. These involve stronger selection on the favorable mutations, as described in the Material and Methods.

**Table 4:** Observed and predicted values of autosomal neutral diversity for a 70 gene region, relative to the values without hitchhiking effects.

| **Xover Rate** | **Observed** | **Integral, NC** | **Sum., NC** | **Integral, C** | **Sum., C** |
| --- | --- | --- | --- | --- | --- |
| **No g.c.** |  |  |  |  |  |
| 0.5 | 0.516 (0.500,0.528)<br>0.430 (0.419,0.441) | 0.582<br>0.487<br>0.461 | 0.612<br>0.530<br>0.479 | 0.471<br>0.450<br>0.409 | 0.507<br>0.469<br>0.432 |
| 1.0 | 0.655 (0.637,0.671)<br>0.555 (0.536,0.573) | 0.713<br>0.597<br>0.592 | 0.733<br>0.610<br>0.606 | 0.633<br>0.545<br>0.534 | 0.659<br>0.562<br>0.553 |
| 1.5 | 0.735 (0.727,0.743)<br>0.631 (0.621,0.643) | 0.786<br>0.669<br>0.664 | 0.801<br>0.680<br>0.676 | 0.726<br>0.620<br>0.621 | 0.745<br>0.633<br>0.635 |
| 2.0 | 0.772 (0.763,0.781)<br>0.675 (0.666,0.683) | 0.832<br>0.716<br>0.712 | 0.843<br>0.724<br>0.721 | 0.784<br>0.676<br>0.676 | 0.799<br>0.687<br>0.688 |
| 2.5 | 0.820 (0.812,0.828)<br>0.715 (0.706,0.724) | 0.862<br>0.744<br>0.740 | 0.871<br>0.750<br>0.748 | 0.823<br>0.711<br>0.711 | 0.835<br>0.720<br>0.720 |
| **G.c.** |  |  |  |  |  |
| 0.5 | 0.685 (0.674,0.695)<br>0.544 (0.534,0.552) | 0.753<br>0.585<br>0.578 | 0.740<br>0.582<br>0.575 | 0.684<br>0.554<br>0.539 | 0.668<br>0.550<br>0.535 |
| 1.0 | 0.767 (0.763,0.771)<br>0.648 (0.626,0.660) | 0.821<br>0.684<br>0.682 | 0.813<br>0.681<br>0.679 | 0.771<br>0.643<br>0.643 | 0.761<br>0.638<br>0.639 |
| 1.5 | 0.815 (0.809,0.821)<br>0.703 (0.690,0.717) | 0.858<br>0.731<br>0.729 | 0.855<br>0.729<br>0.728 | 0.818<br>0.697<br>0.697 | 0.814<br>0.695<br>0.696 |
| 2.0 | 0.850 (0.834,0.856)<br>0.724 (0.713,0.736) | 0.882<br>0.749<br>0.748 | 0.881<br>0.749<br>0.748 | 0.849<br>0.721<br>0.721 | 0.847<br>0.721<br>0.721 |
| 2.5 | 0.863 (0.858,0.869)<br>0.753 (0.744,0.761) | 0.899<br>0.774<br>0.773 | 0.899<br>0.774<br>0.773 | 0.871<br>0.750<br>0.750 | 0.871<br>0.750<br>0.751 |

The entries for the observed values are the mean synonymous site diversities from the simulations with 70 genes, relative to the corresponding values in the absence of selection at linked sites. The upper and lower entries in each cell are the values with SSWs alone and with SSWs and BGS, respectively.
The upper entries in each cell for the predictions are the reductions with SSWs alone; the middle entries use only the BGS effects estimated from neutral sites ( $B_1$ ); the lowest entries also include the BGS effects on adaptive substitution rates obtained from the simulations ( $B_2$ ).
The columns labelled 'Integral' use the approximate integral formulae for SSW effects (equations S24-33); those labelled 'Sum.' use the summation formulae, Equations 5 and 6.
'NC' denotes predictions without correcting for sweep duration (Equation 5). 'C' denotes predictions that correct for sweep duration (Equations 12). 'No g.c.' and 'G.c.' refer to results without gene conversion and with the standard gene conversion parameters, respectively. For the summation predictions, corrections for multiple recombination events during the sweep (Equation S20) and for the variance in sweep time (File S1, section 3) were applied.

Concerning the first point, diversities are considerably overpredicted by the uncorrected values from Equation 5 (but which included the correction for interference), by up to 20% with the lowest rate of crossing over used in Tables 4 and S6. This shows that treating recurrent sweeps as point events can produce significant errors. For the second point, the agreement between the integral and summation results is surprisingly good overall. The largest discrepancies occur when the rate of crossing over is low, and gene conversion and BGS are absent, when they are of the order of 7.6% of the lower value.

For the third point, the agreement between the simulation means and the predictions with the corrections is generally very good, although the integral results underpredict diversity by about 20% for the autosomal case with no gene conversion or BGS. If the correction for interference is not applied, lower diversities are predicted, which sometimes give better agreement with the simulation results, but the effects are not major (Table S7). The main contribution to the improvements in fit from the other corrections comes from the sweep duration, as can be seen from results where one or both of the other factors (multiple recombination events and coalescence during a sweep), as well as interference, are omitted (Table S7). Omission of the correction for coalescence during sweeps usually has the next largest effect, mainly because it reduces the contribution to coalescent time from samples taken during sweeps (section S11 of File S1). Overall, omission of all the corrections except that for sweep duration produces remarkably good results.

With respect to the fourth point above, the fits with *B*_1_alone are good, except for the lowest rate of crossing over and no gene conversion (an error of 9% in Table 4). Overall, it seems that relatively little is to be gained by using *B*_2_.

The predictions of the effects of selective sweeps use a single gene model, based on the assumption that the effects of sweeps with the parameters assumed here are localized to single gene regions. The simulation results with sweeps alone in regions with crossing over (File S2) show that there is no noticeable effect of the numbers of genes on the mean synonymous site diversities, consistent with this assumption. This is not surprising, given that the expected reduction in diversity at a neutral site due to a single sweep at recombination distance *r* is approximately *γ* ^−4*r*/*s*^, where *γ* and *s* are the scaled and absolute selection coefficients for the favorable allele. With the values of *γ* and *s* for autosomal NS mutations assumed here (250 and 1 x 10^-4^ for natural populations, respectively), an effective crossing over rate of 1 x 10^-8^ and a distance of 2000bp between sites (the minimum for sites in separate genes), the expected reduction in diversity with no gene conversion is 250 ^(–0.8)^ = 0.01, which is essentially trivial.

This conclusion does not apply in the absence of recombination, which has been studied theoretically by Kim and Stephan (2003) and Weissman and Hallatschek (2014). In this case, the simulation results displayed in File S2 show that there is a large effect of the number of genes. With no crossing over, gene conversion or BGS, the mean autosomal diversities relative to neutral expectation were 0.0819, 0.0700 and 0.0675 for 70, 140 and 210 genes, respectively. These results can be compared to the predictions from the approximate Equation 5 of Weissman and Hallatschek (2014), modified for diploidy with semi-dominance, which gives the absolute neutral nucleotide diversity under recurrent sweeps with recurrent sweeps as 8μln[2ln(*γ*)/*U_b_*]/*s*. The resulting predicted values are 0.195, 0.183 and 0.176, respectively.

As was also found by Weissman and Hallatschek (2014), the theoretical results thus considerably overpredict diversity. Gene conversion greatly reduces the effects of sweeps, with relative diversities of 0.130, 0.090 and 0.0832 in the absence of BGS. BGS has a much greater effect on diversity than sweeps when crossing over is absent. With gene conversion, it gives relative diversity values of 0.0867, 0.0429 and 0.0293 for 70, 140 and 210 genes, respectively. Essentially the same values are seen with both BGS and SSWs, reflecting the fact that the rate of sweeps is greatly reduced in the presence of BGS (see Table 3). The predicted relative diversity value for a 70 gene region is quite close that observed for the fourth chromosome of *D. melanogaster*, which has a similar number of genes (Campos *et al.* 2014), suggesting that diversity in non-crossover regions of the genome is strongly influenced by BGS, as was also inferred by Hough *et al.* (2017) for the case of the newly evolved Y chromosome of *Rumex*.

## Discussion

### Accuracy of the approximations for pairwise diversity with hitchhiking

We have developed a new expression for the effect of a single substitution of a favorable allele on pairwise neutral diversity at a linked site (Equation 4). This uses an approximate formula for the duration of an adaptive substitution, which includes stochastic contributions (Equations 2 and 3). In addition, we have developed expressions for the effects of coalescence and multiple recombination events during a substitution (sections S3 and S4 of File S1), as well as a crude correction for interference among selective sweeps. More importantly, we have derived new formulae to predict the effects of a constant rate of recurrent adaptive substitutions on pairwise neutral diversity (Equations 12). This approach, while admittedly somewhat heuristic, avoids the assumption made in most previous models of recurrent sweeps that the duration of an adaptive substitution can be neglected, enshrined in Equation 7 (Kaplan *et al.* 1989; Wiehe and Stephan 1993; Kim and Stephan 2000). This equation has been used several times for inferences about sweep parameters (Sella *et al.* 2009; Elyashiv *et al.* 2016; Campos *et al.* 2017), but overestimates diversities compared with the simulations, especially with high rates of adaptive substitutions and low rates of crossing over. The comparisons of the simulation results with the theoretical predictions (Table S7) suggest that the corrections for coalescence and multiple recombination events during a sweep are sufficiently small that they can be ignored for most purposes.

As described at the end of the Results section, the integral approximations provides results that are quite close to the more exact results from summations. Similarly, the use of the reduction in diversity at neutral sites caused by BGS (*B*_1_) for predicting the reduction in rates of substitution of adaptive mutations performs nearly as well as the use of the factor *B*_2_ derived from the simulations. This suggests that inference methods can be simplified by using the integral approximations for both selective sweeps and BGS.

Another feature of the work presented here is the inclusion of gene conversion into sweep models, as was also done by Campos *et al.* (2017) but which has been ignored in previous treatments of sweeps. Gene conversion events that are not associated with crossovers are known to be a major source of recombination events at the intragenic level in *Drosophila* (Hilliker and Chovnick 1981; Hughes *et al.* 2018). With the standard autosomal effective crossing over rate for *D. melanogaster* of 1 x 10^−8^ per bp (Campos *et al.* 2014), the effective rate of crossing over between two sites separated by 500bp is 5 x 10^−6^. With scaled and absolute selection coefficients for NS mutations of *γ* = 250 and *s* = 10^−4^, used for Table 4, the expected proportional reduction in diversity at the end of a sweep for a neutral site that is 500bp away from the selected site is approximately *γ* ^(–4*r/s*)^ = 250 ^(–4x 0.05)^ = 0.33. With the somewhat conservative gene conversion parameters assumed in the table, Equation 1 of Frisse et al. (2001) implies an additional contribution to the effective recombination rate of 4.4 x 10^-6^, so that the total effective recombination rate is 9.4 x 10^-6^. This yields a reduction in diversity of 0.13, approximately 40% of the value with no gene conversion. Consistent with this result, the simulation results and theoretical predictions are significantly affected by gene conversion, such that the expected effects of sweeps on diversity are considerably reduced if gene conversion is present (Tables 4 and S6). Ignoring gene conversion in sweep models is likely to substantially bias estimates of sweep parameters.

While our models assume ‘hard’ sweeps, where the new favorable mutation is introduced as a single copy, Equation 12a can also be applied to other situations, such as ‘soft’ sweeps arising from standing variation or multiple mutations to the favorable allele at a locus (Hermisson and Pennings 2005, 2017). The only modification that need be made is to the expression for the reduction in diversity immediately after a sweep (*D*) in Equation 8.

### Interference between adaptive substitutions

Our results suggest that there is a minor, but noticeable, degree of interference among adaptive substitutions in the presence of crossing over (File S1, section S8). A somewhat counter-intuitive finding is that the proportion of substitutions eliminated by interference is nearly independent of the strength of selection and the recombination rate. Relatively weak effects of the recombination rate, provided it is not very high or low, are also evident in Table 3 of Kim and Stephan (2006) and in Figure 3 of Barton (1995), for cases when the selection coefficients at the selected loci are similar. A possible explanation for this is that recombination has a dual effect on the potential for interference. If a second favorable mutation arises early enough during the substitution of a previous one, it is likely to be on the wild-type background, so that a recombination event that puts it onto the mutant background would enhance its fixation probability. The opposite would be the case if it arises late during the sweep. Similarly, the faster (lower) rate of spread of a more strongly (weakly) selected mutation would reduce (increase) the probability of either type of recombination event, so that its selective advantage might not greatly affect the opportunity for interference. Finally, the product of the rate of substitution of favorable mutations and the time taken for a substitution (*ωT_s_*) determines the chance of two substitutions overlapping in time. Because *ω* is proportional to the scaled selection coefficient *γ* and *T_s_* is close to being inversely proportional to *γ* (Equation 4), *ωT_s_* is only weakly dependent on *γ*.

### The relation between sequence diversity and rate of crossing over

It is also of interest to ask what light the theoretical results described above shed on the observed positive relationship between DNA sequence variability at putatively neutral or nearly neutral sites within a gene in *D. melanogaster* and the local rate of recombination experienced by the gene (Aguadé *et al.* 1989; Begun and Aquadro 1992). This observation stimulated interest in models of SSWs and BGS, and its possible causes have been a long-standing subject of debate; for reviews, see Sella *et al.* (2009), Stephan (2010), Cutter and Payseur (2013) and Charlesworth and Campos (2014).

Recent analyses of population genomic data have confirmed the existence of a strong relationship between synonymous nucleotide site diversity (*π*_*S*_) and the effective rate of crossing over, even after excluding genes in non-crossover regions. For example, Figure 2 of Charlesworth and Campos (2014) shows the regressions of *π*_*S*_for autosomal and X-linked genes on their effective rates of crossing over, for a sample from a Rwandan population of *D. melanogaster*. The ratios of the value of *π*_*S*_for an effective rate of crossing over of 0.5 cM/Mb to the value with a crossing over rate of 2 cM/Mb (the upper limit to the autosomal rate) are *K* = 2.38 for autosomes and *K* = 1.78 for the X chromosome. The simulation results in Table 4 for both BGS and SSWs with 70 autosomal genes and gene conversion give *K* = 1.33 and *K* = 1.24 with SSWs alone; *K* = 1.27 with BGS alone (Table 3), a much weaker relationship than is observed.

What causes this discrepancy? One possibility is that the mean scaled selection coefficients for favorable mutations used in these simulations are unrealistic. This was checked by re-running the calculations with different *γ* values for the favorable mutations, using the summation method with all the corrections, and *B*_1_ for the effects of BGS, as well as selection, mutation and recombination parameter values appropriate for the natural population rather than the simulations (see Table 1). For the reasons given in the final part of the Discussion, we used a linear genetic map and no correction for the repulsion BGS terms when determining the values of *B*_1_, with selection, mutation and recombination parameter values. We also used the rate of gene conversion indicated by experimental studies in *D. melanogaster* (Hilliker *et al.* 1981, 1994; Miller *et al.* 2016), instead of the conservative value used previously (half of the empirical value). Because the effect of interference among sweeps is unknown for these parameters, it was found to be small in the presence of gene conversion and BGS, it has been ignored in the following analyses.

Becaues *B*_1_ is somewhat sensitive to the number of genes modeled, we used results for 210 genes, approximating the behavior of a small group of linked genes with similar recombination rates. In the absence of selective sweeps, but with gene conversion and BGS, *K* = 1.17 for the autosomal model, reflecting the greater effect of recombination with a linear map than with the Haldane mapping function. If the standard *g* values for favorable mutations are used for the sweep predictions, and BGS and gene conversion are included, *K* = 1.27, somewhat lower than the simulation result. The ratios for *γ* values that are half and twice the standard values are 1.19 and 1.47, respectively.

There is thus only a rather weak dependence on the strength of selection on favorable mutations. This is not surprising, in view of the fact that the net effect of sweeps on a neutral site for favorable mutations is proportional to *γ* and the product of the deterministic component of Equation 4. Its logarithm is thus approximately equal to a constant plus ln(*γ*)[1 – 4*ρ*γ**^−1^]. For *ρ*γ**^−1^ < < 1, the derivative of this expression with respect to *γ* is approximately equal to *γ*^−1^, which means that there is only a small proportional effect on diversity of a change in *γ* when *γ* >> 1. Similarly, its derivative with respect to *ρ* to is – 4*γ*^−1^ ln(*γ*), which is << 1 when *γ* >> 1. It thus seems unlikely that the weak predicted dependence of neutral diversity on the rate of crossing over can be explained by the choice of selection coefficients for favorable mutations.

The effect of the proportion of mutations that are favorable can be examined in a similar way. With the standard selection coefficients, halving these proportions leads to *K* = 1.24, and doubling them to *K* = 1.34. Although this parameter has a large effect on diversity, its effect is only weakly dependent on the crossing over rate over the range considered here. Even if both the strengths of selection and the proportions of beneficial mutations are doubled, *K* is increased to only 1.64. To explain the observed relation between diversity and rate of crossing, considerably larger values of both the strength of selection and proportion of favorable mutations than are currently suggested by population genomic analyses seem to be required.

Another possibility is that intergenic and intronic sequences are subject to selection, rather than being selectively neutral. Charlesworth (2012b) used evidence on the levels of selective constraints on different types of *Drosophila* DNA sequences to obtain crude estimate of *γ* values for deleterious mutations in weakly constrained and strongly constrained noncoding sequences, as well as for deleterious NS mutations. His analysis showed that a linear genetic map provided a good approximation to the BGS predictions. We used this approach to predict the background selection parameter *B*_1_for a genic region with a given rate of crossing over, modifying it to include the effects of gene conversion, as described in File S1, section S9. For a model of an autosome with the standard rate of gene conversion, this procedure gives *B*_1_values of 0.379, 0.616, 0.724, 0.785 and 0.825 for relative rates of crossing over of 0.5, 1, 1.5, 2 and 2.5, respectively, yielding a ratio 2.07 for relative crossing over rates of 2 and 0.5. If these values are used in the above method for predicting *π/θ* for a natural population with BGS and SSWs, with the standard *γ* values for favorable mutations, the predictions for the different relative rates of crossing over become 0.343, 0.562, 0.652, 0.709 and 0.753, respectively, giving *K* = 2.05.

Both predicted values of *K* are still somewhat lower than the observed value of 2.38. This may reflect the fact that BGS on intergenic sequences is likely to have a weaker effect on the fixation probabilities of favorable mutations than is predicted by *B*_1_when crossing over rates are relatively high, given their distance from positively selected mutations in coding sequences, so that the effect of increased crossing over is greater than predicted by this model. Another possibility is that a class of much more strongly selected favorable mutations (e.g. Sella *et al.* 2009) is contributing to intergenic effects of sweeps that are not captured by single locus models, and which could affect the *K* values.

The same procedure can also be applied to the X chromosome, as described in section S10 of File S1. If the low gene density in low crossing over regions of the *D. melanogaster* X chromosome is taken into account (Campos *et al.* 2014), a moderately good fit to the data is obtained for the model with SSWs and BGS, both for the relation between crossing over rate and diversity on the X, and for the X/autosome diversity ratios at the two extremes of the autosomal crossing over rates. The rates of substitution of favorable NS autosomal and X-linked mutations can be analysed in a similar way, and also show a reasonable level of agreement with the observations (section S11 of File S1).

These analyses are very crude, and require considerable refinement, but suggest that the relative values of nearly neutral variability and rates of adaptive evolution in crossover regions of the *D. melanogaster* genome are more strongly influenced by BGS rather than SSWs, in agreement with Comeron (2014). As discussed at the end of the Results section, this conclusion also applies to regions of the genome with zero or very low rates of crossing over, where the effects of SSWs are expected to be weak.

### Distortion of the site frequency spectrum by hitchhiking

We have not previously discussed the effects of BGS and SSWs on the site frequency spectra (SFS) at the neutral loci affected by selection at linked sites in genomic regions with crossing over. While it should be possible to use the theoretical frameworks developed for BGS (Zeng and Charlesworth 2011; Nicolaisen and Desai 2013) and SSWs (Durrett and Schweinsberg 2004; Kim 2006; Pfaffelhuber *et al*. 2006; Bossert and Pfaffelhuber 2013), this would require extensive calculations that are outside the scope of this paper. We note, however, that the simulation results shown in File S2 show that recurrent SSWs have noticeable effects on the SFS, even with quite high rates of crossing over, in the direction of an excess of rare variants over neutral expectation, as expected from previous theoretical work (Kim 2006), and as has been seen in previous simulation studies, e.g. Messer and Petrov (2013). Such effects of BGS and SSWs on the SFS may bias estimates of demographic parameters when neutrality is assumed (Messer and Petrov 2013; Ewing and Jensen 2016; Schrider *et al.* 2016).

For example, with 70 autosomal genes and gene conversion, the mean values of synonymous site Tajima’s *D* per gene with SSWs and BGS for relative rates of crossing over of 0.5, 1, 1.5, 2.0, and 2.5 were – 0.209, – 0.156, – 0.116, – 0.111 and – 0.069, respectively. The corresponding mean proportions of singletons were 0.319, 0.310, 0.302, 0.299 and 0.295, compared with the neutral value from simulations of 0.275. In the presence of BGS but not SSWs, the mean values of Tajima’s *D* were – 0.046, – 0.013, – 0.019, – 0.036 and 0.000, respectively, compared with the neutral value of 0.042. The mean values of the proportions of singletons were 0.288, 0.286, 0.284, 0.289 and 0.282. Thus, with the parameters used here, BGS contributes very little to the distortion in the SFS, consistent with previous theoretical work on BGS with significant amounts of recombination (Zeng and Charlesworth 2011; Nicolaisen and Desai 2013). Detailed comparisons with the data are made difficult by the probable effects of demographic factors on these measures of distortion of the SFS, which will tend to obscure the effects of selection at linked sites, especially their relations with the rate of crossing over.

As might be expected, stronger selection on favorable mutations increases the extent of distortion of the SFS. For example, with the stronger of the two selection models for the X chromosome, the Tajima’s *D* values and proportions of singletons for the standard rate of crossing over for 70 genes with gene conversion, SSWs and BGS were – 0.434 and 0.360, respectively. The difference between X and autosomes is qualitatively similar to what is seen for the Rwandan population of *D. melanogaster*, shown in Figure 2 of Campos *et al.* (2014). However, the difference between X chromosome and autosomes of the distortion of the SFS is much greater than is seen in the simulations. It remains to be seen whether demographic effects can explain this discrepancy.

The picture is, however, very different when crossing over is absent. For 70 autosomal genes with gene conversion, the means of Tajima’s *D* and the proportion of singletons for synonymous sites with BGS alone were – 0.880 and 0.488, respectively. With SSWs as well, the values were changed by relatively small amounts, to – 1.306 and 0.563, respectively, reflecting the greatly reduced rate of fixations of favorable mutations when crossing over is absent (Table 3). It therefore seems likely that the distorted SFSs seen in genomic regions that lack crossing over (Cutter and Payseur 2013; Campos *et al.* 2014) are mainly caused by BGS in the weak interference selection limit, when interference among sites subject to purifying selection causes genealogies at linked sites to have longer terminal branches relative to neutral expectation (Gordo *et al.* 2002; Kaiser and Charlesworth 2009; Seger *et al.* 2010; O'Fallon *et al.* 2010; Good *et al.* 2014).

### Problems with simulating BGS

We conclude with a discussion of some technical questions concerning the modeling of BGS in SLiM. As described in the first part of the Results section, the fact that SLiM assumes a lack of crossover interference requires the modification of the standard BGS equations to model the Haldane mapping function, as described in the section S1 of File S1. In addition, for accurate approximations to the simulation results, it was necessary to include an additional term in the BGS equations that results from deleterious mutations that were in initially in repulsion with a new neutral variant (Santiago and Caballero 1998; Charlesworth 2012b); this is ignored in the equations that are usually used to model BGS.

These properties are more a reflection of the simulation procedure than of biological reality. Equation S1b implies that the extra term added to the standard BGS equation of Nordborg *et al.* (1996) is proportional to the sum of twice the product of the deleterious mutation rates and the mean of *t* = *hs* for deleterious mutations, multiplied by a term that is nearly independent of the factor used for rescaling. This term is exactly equal to this product when there is no recombination, and is then equal to the additive genetic variance of fitness under deterministic mutation-selection balance (Mukai *et al.* 1972). Since the deterministic parameters that are thought to be realistic for a Drosophila population have been multiplied by 532 for use in the simulations, the additive genetic variance in fitness is multiplied by a factor of (532)^2^ = 283,024 compared with its value for the real population. With 70 genes, for example, the additive variance in the simulations is 0.0542, whereas the corresponding value for the population is 1.92 x 10^−7^. In contrast, the Nordborg *et al.* (1996) equation depends largely on the ratios of deterministic parameters, except for the multiplication of the recombination rate by a factor of 1 – *t*, and so is largely unaffected by the rescaling. In the real population, this additional term is effectively negligible, justifying the use of the standard equation for modeling BGS, e.g. (McVicker *et al.* 2009; Charlesworth 2012b; Comeron 2014; Elyashiv *et al.* 2016; Campos *et al.* 2017).

The use of the Haldane mapping function also means that the simulated rate of recombination for the region as a whole is affected by the rescaling, since the frequency of double crossovers is greatly increased over what would be found in a region of the same physical length in the real population. For example, with the standard rate of crossing over and 70 genes, the map length of the region with the standard rate of crossing over is 1.62. With a Poisson distribution of numbers of crossovers, as assumed in the simulations, the proportion of double crossovers among chromosomes that have experienced a crossover is 0.5 x (1.62)^2^ x exp(– 1.62)/[1 – exp(– 1.62)] = 0.324. For regions of the size that we have simulated, the high level of crossover interference in *Drosophila* (Hughes *et al.* 2018) means that a linear relation between the frequency of crossing over and physical distance is close to reality for a real population (Charlesworth 2012b). Unfortunately, except for the cases with a frequency of crossing over of one-half the standard rate used here, it is impossible to simulate a linear model with 70 genes or more, since the expected number of crossovers in the region is greater than one, which is inconsistent with a model that assumes that there is either a crossover or no crossover in the region.

Given that our simulation results generally support the use of the theoretical formulae for both background selection and selective sweeps, largely because both BGS and SSW effects extend over much smaller distances than the whole region, this implies that the use of formulae based on the BGS and SSW equations with a linear genetic map is probably justified for most analyses of population genomic data, although it would be desirable to validate this conclusion with simulations using much larger population sizes than was feasible here.

## Acknowledgments

This work was supported by grant RPG-2015-2033 from the Leverhulme Trust (to BC). We thank Hannes Becher, Graham Coop, Matty Hartfield and Lei Zhao for useful discussions and comments on the manuscript, and Jeff Jensen, Peter Keightley and two anonymous reviewers for their comments on the manuscript.

